# A peptide pair coordinates regular ovule initiation patterns with seed number and fruit size

**DOI:** 10.1101/736439

**Authors:** Nozomi Kawamoto, Dunia Pino Del Carpio, Alexander Hofmann, Yoko Mizuta, Daisuke Kurihara, Tetsuya Higashiyama, Naoyuki Uchida, Keiko U. Torii, Lucia Colombo, Georg Groth, Rüdiger Simon

**Affiliations:** Institute for Developmental Genetics, Heinrich-Heine University Düsseldorf, University Street 1, D-40225 Düsseldorf, Germany; Cluster of Excellence on Plant Sciences (CEPLAS); Agriculture Research division, Agriculture Victoria, Australia; Institute of Biochemical Plant Physiology, Heinrich-Heine University Düsseldorf, University Street 1, D-40225 Düsseldorf, Germany; Institute for Advanced Research (IAR), Nagoya University, Furo-cho, Chikusa-ku, Nagoya, Aichi 464-8601, Japan; Institute of Transformative Bio-Molecules (ITbM), Nagoya University, Furo-cho, Chikusa-ku, Nagoya, Aichi 464-8601, Japan; JST, PRESTO, Furo-cho, Chikusa-ku, Nagoya, Aichi 464-8601, Japan; Department of Biology, University of Washington, Seattle, WA, 98195 USA; Howard Hughes Medical Institute and Department of Molecular Biosciences, University of Texas at Austin, Austin, TX, 78712 USA; Universita degli studi di Milano, Italy

## Abstract

Ovule development in *Arabidopsis thaliana* involves pattern formation which ensures that ovules are regularly arranged in the pistils to reduce competition for nutrients and space. Mechanisms underlying pattern formation in plants, such as phyllotaxis, flower morphogenesis or lateral root initiation, have been extensively studied, and genes controlling the initiation of ovules have been identified. However, how a regular spacing of ovules is achieved is not known. Using natural variation analysis combined with quantitative trait locus analysis, we found that the spacing of ovules in the developing fruits is controlled by two secreted peptides, EPFL2 and EPFL9 (also known as Stomagen), and their receptors from the ERECTA (ER) family that act from the carpel wall and the placental tissue. We found that a signalling pathway controlled by EPFL9 acting from the carpel wall through the LRR-receptor kinases ER, ERL1 and ERL2 promotes fruit growth. Regular spacing of ovules depends on EPFL2 expression in the carpel wall and in the inter-ovule spaces, where it acts through ERL1 and ERL2. Loss of EPFL2 signalling results in shorter fruits and irregular spacing of ovules or even ovule twinning. The EPFL2 expression pattern between ovules is under negative-feedback regulation by auxin, which accumulates in the arising ovule primordia. We propose that the auxin-EPFL2 signalling module evolved to control the initiation and regular, equidistant spacing of ovule primordia, which serves to minimise competition between developing seeds. Together, EPFL2 and EPFL9 coordinate ovule patterning and thereby seed number with fruit growth through a set of shared receptors.

## Introduction

Plants have evolved diverse strategies to maximise their reproductive success, which enables them to transfer genetic resources to subsequent generations. To produce floral organs at an appropriate time, plants integrate various environmental cues to induce flowering^1, 2^. When this process is triggered in *Arabidopsis thaliana*, each flower produces four sepals, four petals, six stamens and one pistil which originates from the fusion of two carpels. The ovules, which contain the egg cells, reside in the pistil and are derived from another meristematic tissue within the pistil termed placenta^2, 3^, where they are almost simultaneously initiated in two parallel rows within each carpel. The number of ovules per flower determines the maximum number of seeds that a single flower can generate. At the transition from flower developmental stage 8 to 9 as defined by Smyth et al.^4^, ovules are initiated from the placenta with regular 2 to 4 cell intervals. This regularity enables plants to reduce competition between adjacent ovules or the developing seeds after fertilization; disruption of this regular pattern could result in the formation of small or large, or closely juxtaposed ovules, which would bias reproductive success depending on random positional effects. During further growth, the pistil forms a silique that encloses the developing seeds until they reach maturity and are shed. Thus, the overall size of the silique places a natural constraint on the number of seeds that can be formed, their final size, or both, and silique growth needs to be tightly coordinated with the ovule initiation process. Indeed, final fruit length is normally well correlated with the number of seeds grown within it^5^. However, how this coordination is achieved, and how ovules are initiated strictly at very regular intervals remains to be investigated.

Successful ovule formation is by itself a prerequisite for seed production and has therefore attracted widespread research interest, so that mechanistically, the process of ovule primordia formation is at least partially understood and the functions of key regulatory genes have been identified^2^. Ovule primordia originate from periclinal divisions in subepidermal cell layers of the placenta, and their formation requires the coordinated activity of auxin and cytokinin signalling pathways. Here, PIN1 acts as the main auxin transporter, and *pin1-5* mutants develop pistils with a reduced ovule number^6^. The expression of *PIN1* is further modulated by cytokinin. An increase in cytokinin levels due to loss of cytokinin degrading enzymes causes an increase in ovule number per flower, possibly by upregulation of PIN1 levels^7^. Other phytohormones involved are Gibberellins (GAs) and Brassinolide (BR), which act antagonistically to restrict (GA) or promote (BR) ovule formation via regulation of cytokinin signalling^8, 9^. In response to auxin, the transcription factor MONOPTEROS/AUXIN RESPONSE FACTOR5 (MP/ARF5) is activated and regulates the expression of the transcription factors AINTEGUMENTA (ANT), CUP SHAPED COTYLEDON1 (CUC1) and CUC2 in ovule primordia and the boundary domains between ovules, respectively^10^. A knockdown of *CUC1* expression in *cuc2* or *cuc2;ant* mutant backgrounds reduces ovule numbers, whereas *cuc2;cuc3* double mutants give rise to fused ovules, indicating that the generation of interorgan boundaries depends on partially overlapping CUC functions. CUC1 and CUC2 affect *PIN1* expression via control of cytokinin inactivating enzymes^7^. Overall, the process of ovule formation strongly resembles that of other lateral organs, where initials are first defined by local auxin accumulation^10^. The distance between ovule primordia determines ultimately the total number of seeds that can be generated on a single flower, if silique length is constant. A recent genome wide association study identified *NEW ENHANCER OF ROOT DWARFISM1* (*NERD1*) as a positive regulator of ovule number, however, the *nerd1* mutants generated drastically shortened siliques, suggesting that *NERD1* does not play a specific role in controlling the distance between arising ovules^11^. Alonso-Blanco and colleagues previously proposed the *ERECTA* (*ER*) locus of Arabidopsis to be a major determinant of several life history traits, among them fruit size and ovule number per flower^5^. The *er* mutants are characterised by a short-fruit phenotype and a compact shoot architecture in the Landsberg background^12^. The *ER* gene encodes a leucine rich repeat (LRR) receptor kinase which regulates pattern formation in multiple developmental pathways, including stomata development, vascular architecture and leaf margin serration^13^; the related ER-family genes *ERECTA-LIKE1* (*ERL1*) and *ERL2* contribute partially overlapping functions with ER^14–17^. Ligands for ER-family receptors belong to the evolutionary conserved EPIDERMAL PATTERNING FACTOR (EPF)/EPF-LIKE (EPFL)-family of cysteine-rich secreted peptides, with 11 members in Arabidopsis^18^. Some EPF/EPFL peptides act antagonistically in stomata development by competing for interaction with the receptor complex, and consequently trigger different signalling readouts in the stomata lineage^18–26^. For example, while EPF2 activates the MAPK cascade upon binding to the ER/ERL1/TMM receptor complex to restrict entry of epidermal cells into the stomatal lineage, EPFL9/STOMAGEN competes for binding and interacts preferentially with ER/ERL1^25^. Because EPFL9 binding does not induce MAPK activation, SPCH is not degraded, resulting in the production of supernumerous stomata^27, 28^. Beyond epidermal cell specification, EPFL2 was found to interact with ER, ERL1 and ERL2 to promote leaf margin tooth growth via regulation of auxin responses^17^.

We started to investigate the underlying mechanisms of regular ovule initiation by asking whether the spacing of ovules is largely genetically or environmentally controlled. A natural variation analysis combined with QTL analysis of candidate lines first identified *ER* and *EPFL2* as key loci that control the spacing of ovules in the developing fruit. Detailed genetic and gene function analyses further revealed that at least two separate pathways involving members of the ER and EPF families control regular spacing of ovules, together with fruit size. We propose that the tight coupling of fruit growth with ovule initiation at regularly spaced intervals depends on the negative feedback regulation between EPFL2 and auxin. The output of this patterning module would safeguard a low variance between ovules as a conservative bet hedging approach.

## Results

### ER regulates the density of ovules

Because key genes controlling ovule formation such as *PIN1*, *MP* or *CUC1* act in multiple processes of organogenesis in plants, their interactions might be hardwired, and classical mutant screens might not deliver insights into the regulation of ovule density patterning itself. We therefore first resorted to study the range of variation in ovule density that can be observed between natural accessions of Arabidopsis. We grew the accessions at two different temperatures as a proxy to further access environmental control of this patterning process. For 96 accessions, we measured fruit length and the number of seeds including unfertilized ovules (= total number of ovules) per fruit in stage 17 flowers (all stages according to Smyth et al.,^4^), and calculated total ovule density as a derived trait (total number of ovules per mm fruit length). The total ovule density strongly varied between accessions and temperatures, ranging from 2.37 to 6.36 (N/mm) (Figure S1). Our natural variation analysis revealed several accessions with a characteristic ovule-density phenotype (Figure 1A, B), which was only mildly affected by temperature. Hence, we sought regulators by applying QTL analysis to L. *er* x Cvi-0 recombinant inbred lines (RILs), since Cvi-0 has long fruits and a low ovule density, whereas L. *er* carries shorter fruits with a high ovule density (Figure 1A, B). QTL analysis allowed us to find a significant peak on chromosome 2 (Figure 1C). Among many loci in this chromosomal region, the *ER* locus seemed to be the most influential candidate. The accessions Landsberg *erecta* (L. *er*) and Vancouver-0 (Van-0) were characterized by shorter fruit and a higher ovule density compared to other accessions (Figure 1B, Figure S1). The L. *er* and Van-0 accessions both carry mutations in the *ER* locus and are known *er* loss-of-function mutants^29, 30^. In addition to these two accessions, Hiroshima-1 (Hir-1) is also known as an *er* loss-of-function mutant^29, 30^. To test the functional importance of *ER* gene in the control of ovule density, we assessed fruit phenotypes in the *er* mutant lines complemented with a wildtype copy of the *ER* locus^29–31^. The short fruit and high ovule density phenotypes in all three accessions were complemented by functional *ER* genomic DNA from Columbia (Col) (Figure S2). Furthermore, the *er-105* mutant in a Col background showed a similar phenotype to the L.*er*, Van-0 and Hir-1 accessions (Figure 2A, Figure S2, S3). These results clearly indicate that *ER* is necessary for the control of ovule density, and that ER acts similarly in different genetic backgrounds.

**Figure 1.**
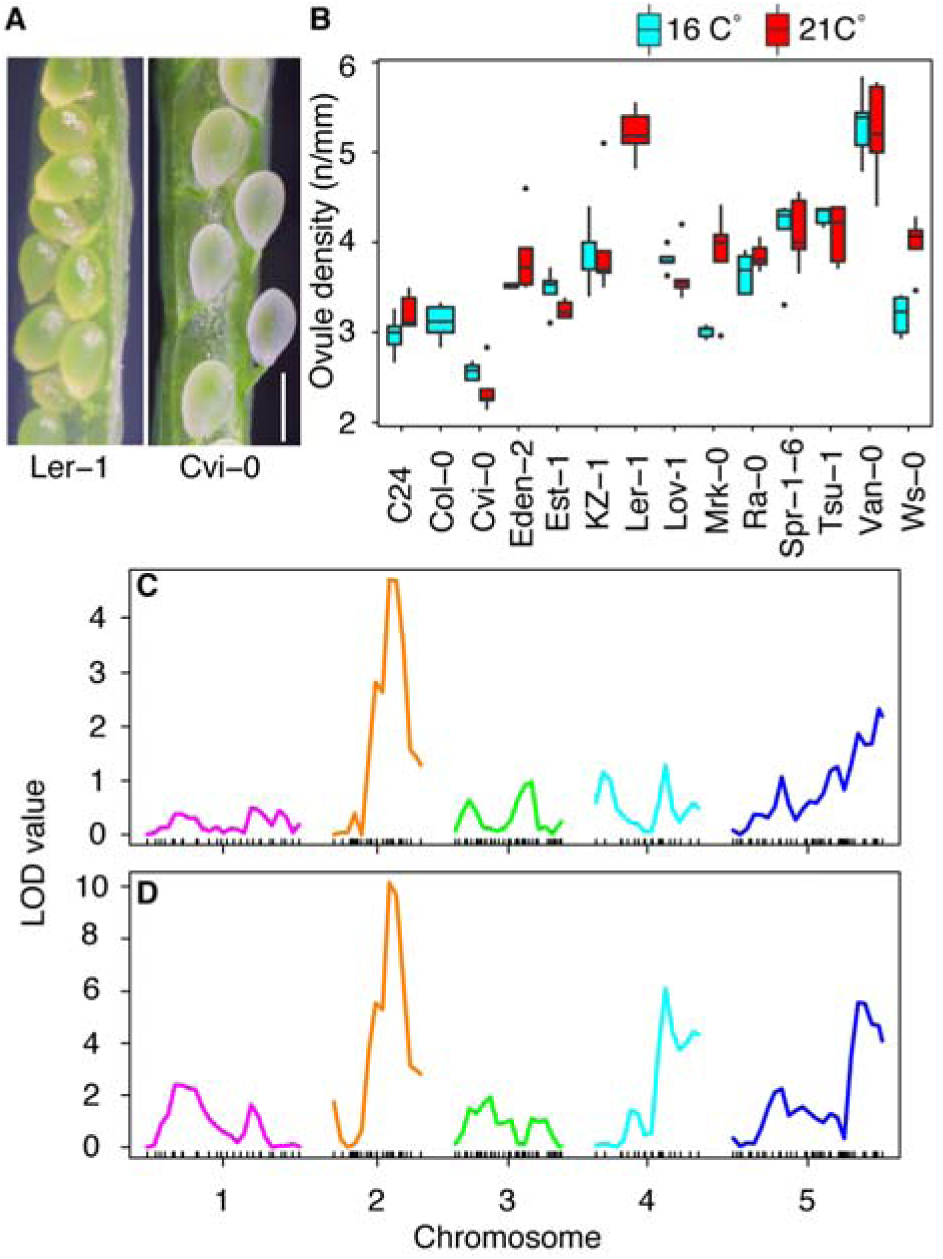
Identification of responsible loci for the reproductive traits. (A) Image of seed density in L. *er* (left) and Cvi-0 (right). Bar = 1 mm. (B) Natural variation analysis on seed density (seed number/fruit length (mm)) phenotype at 16°C (cyan) and 21°C (red). Fourteen selected representative accessions are presented. See the supplementary figure 1 for the phenotype of all accessions. (C) QTL analysis using L. *er* x Cvi-0 recombinant inbred lines [45]. (D) QTL re-analysis with ER as a cofactor. X and Y axes indicate chromosome position and LOD values, respectively.

**Figure 2.**
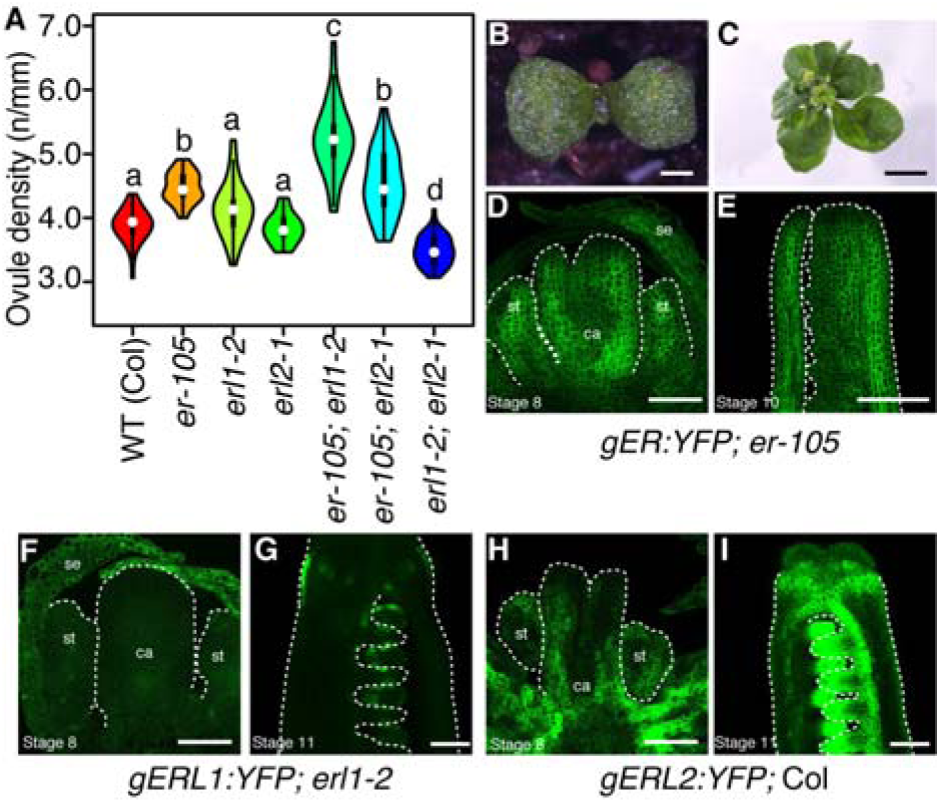
Genetic and expression analysis of ER family receptors. (A) Seed density (seed number/fruit length (mm)). 40 fruits were measured from 3 plants in each genotype. (B) Two weeks old *er-105; erl1-2; erl2-1* plant. (C) Six weeks old *er-105; erl1-2; erl2-1* plant. Expression patterns of ER, ERL1 and ERL2 in stage 8 flower (D, F, H) or later stage (E, G, I) of developing pistils. se, st, ca, v and r indicate sepals, stamens, carpels, valve, replum, respectively. (D, F, H) and (E, G, I) share same scales, respectively. Scale bar = 1 mm (B), 5 mm (C) and 50 µm (D, E, F, G, H, I). Tukey-Kramer’s test was used for the statistical analysis. Different letters indicate significant difference (p< 0.005).

### ERL1 and ERL2 antagonistically function to ER in the regulation of ovule density

Two *ER* paralogous genes, *ERL1* and *ERL2* have overlapping, yet distinct functions with *ER* in the regulation of plant architecture^11^. To investigate the potential role of *ERL1* and *ERL2*, we analyzed *ER*-family receptor mutants in a Col background^15^. Although *erl1-2* and *erl2-1* single mutants did not display an obvious phenotype, the *erl1-2*; *erl2-1* double mutant developed shorter fruits and lower ovule density than the wild-type (Figure 2A, Figure S3). This is in contrast to *er*-mutants, which carried also shorter fruits, but with a higher ovule density than wild type. When combined with *er-105*, either *erl1-2* or *erl2-1* further enhanced the fruit length phenotype and displayed reduced total ovule number, but surprisingly an even higher ovule density (Figure 2A, Figure S3). Since *er-105; erl1-2; erl2-1* triple mutants are very dwarfed and do not produce proper flower organs^32^ (Figure 2B, C), it was impossible to analyze their fruit or ovule density phenotype. However, our results indicate that *ERL1* and *ERL2* function jointly with *ER* to promote fruit growth, whereas two separable *ERL1/2* and *ER* dependent pathways antagonistically regulate ovule density.

### EPFL9 is a ligand for ER family receptors that controls fruit elongation

Our genetic analysis suggested that two independent pathways antagonistically control ovule density- *ER* functions to decrease ovule density whereas *ERL1* with *ERL2* increase ovule density. This suggests that unknown ligands, binding to ER-family receptors, may be involved in ovule density control. Previous work identified EPF/EPFL family peptides as ligands of ER-family receptors that control a variety of biological processes^19, 20, 21–26, 33, 34, 35^. We found that within the *EPF*/*EPFL* family, *EPFL9* is expressed in developing fruits by searching a public expression database. EPFL9 has a unique function in stomatal patterning^21–23^, since all other EPF/EPFL peptides except EPFL9 reduce the number of stomata by activating a downstream MAPK cascade via ER family receptors^17, 18, 19–26, 34, 35^. EPFL9 also interacts with ER-family receptors, but its binding does not activate a MAPK cascade^25^, thus acting in an antagonistic manner to the other EPF/EPFL peptides. We then hypothesized that EPFL9 might function as a ligand for the control of ovule density. Since an EPFL9 mutant was not available, we analyzed STOMAGEN RNAi plants^21^ and found a clear reduction of fruit length and a higher ovule density than in wild-type plants (Figure 3A, Figure S4). Together, the phenotype was weaker than that of *er-105* mutants (Figure 2A, Figure S3). Our result suggests that EPFL9 functions through ER family receptors to promote fruit growth, possibly in conjunction with other related ligands.

**Figure 3.**
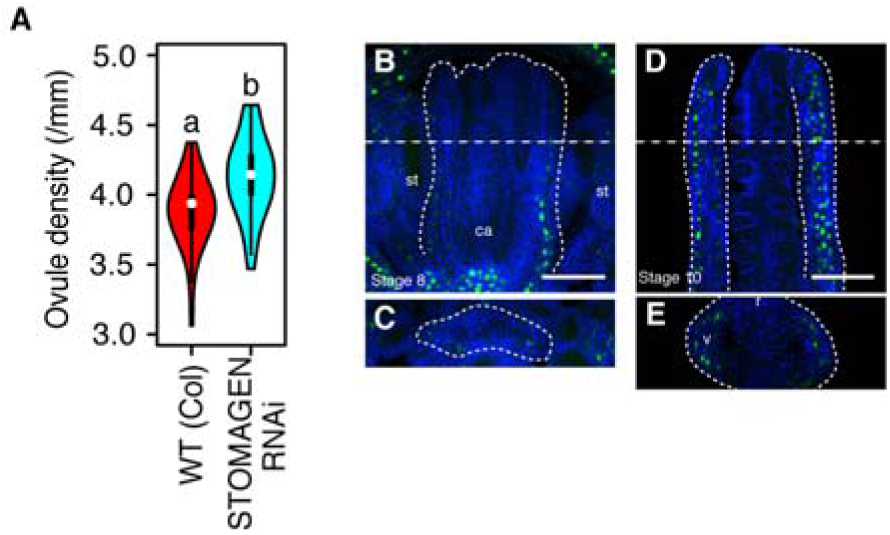
Identification of EPFL9 as a potential ligand for ER. (A) Seed density (seed number/fruit length (mm)). 40 fruits were measured from 3 plants in each genotype. (B-E) Expression pattern of EPFL9 at stage 8 flower (B, C) and stage 9 flower (D, E). Transverse cross section (C, E) were obtained along the lines in (B) and (D). st, ca, v and r indicate sepals, stamens, carpels, valve, replum, respectively. Histone H2B fused EGFP was used as a reporter. Bar = 50 µm. Student’s t-test was used for the statistical analysis. Different letters indicate significant difference (p< 0.005).

### EPFL2 as a ligand for ER-family receptors in ovule spacing

As we described above, EPFL9 functions as a ligand of ER in the stomata pathway. We sought further regulators of ovule density by re-analyzing our QTL data set using the ER marker on chromosome 2 as a cofactor. Cofactor analysis allowed us to improve the detection power and decrease a type II error (false negative)^36^. QTL analysis revealed additional contributing regions on chromosome 4 and 5 for the control of seed density (Figure 1D). Among the candidate loci, we focused on *EPFL2* (At4G37810) on chromosome 4, which acts with ER-family receptors to control leaf serration^17^. The *epfl2-1* mutation in the L.*er* background caused, compared to L.*er*, a minor fruit shortening but a major reduction in ovule number, so that the resulting ovule density was lowered (Figure S5). When *ER* genomic DNA was introduced into the *epfl2* L.*er* accession (L.*er ER*+), the phenotype was still characterized by short fruit length, a low ovule number and a low ovule density (Figure 4A, Figure S5), indicating that *EPFL2* acts independently of *ER*. Overall, the *epfl2-1* phenotype closely resembled that of *erl1-2; erl2-1* double mutants (Figure 2A, Figure 4A, Figure S3, Figure S5), suggesting that ERL1 and ERL2, and not ER, are the key receptors for perception of EPFL2. Since *erl1* and *erl2* mutants are in the Col accession, we generated the novel *epfl2-2* mutant allele in the same genetic background using CRISPR/Cas9 for further genetic analysis. We then crossed *epfl2-2* with *erl1-2; erl2-1* to generate *epfl2-2; erl1-2; erl2-1*, and with *er-105* to generate *epfl2-2; er-105*. The *epfl2-2; erl1-2; erl2-1* triple mutant showed a similar phenotype to the parental lines *epfl2-2* and *erl1-2; erl2-1* (Figure 4B, Figure S8). Furthermore, *epfl2-2; er-105* displayed an additive phenotype, as observed in L. *er; epfl2-1* (Figure 4B, Figure S5, Figure S8). We conclude that EPFL2 mainly functions with ERL1 and ERL2, and not with ER.

**Figure 4.**
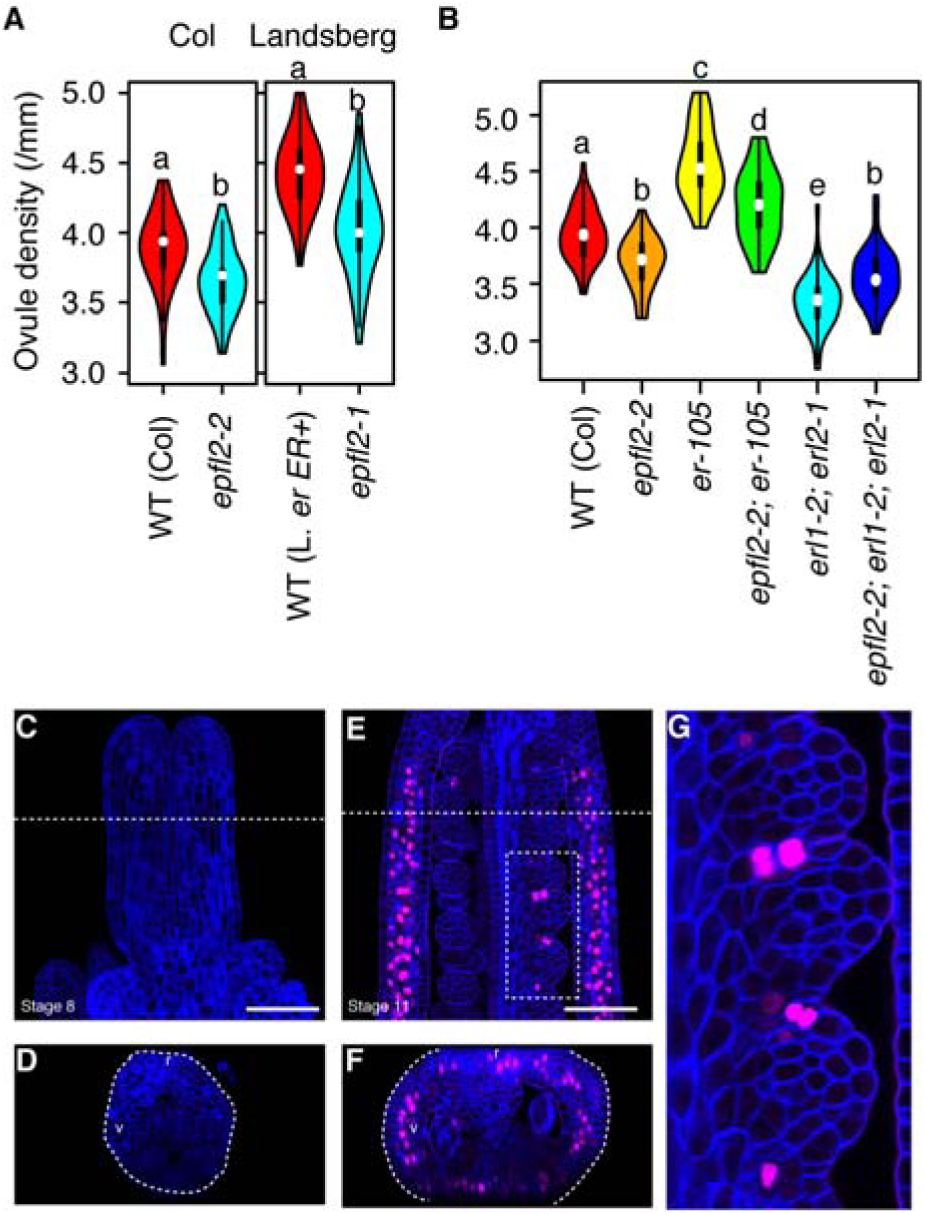
Identification of EPFL2 as a patterning regulator of ovule initiation. (A) Seed density (seed number/fruit length(mm)). 40 fruits were measured from 3 plants in each genotype. (B) Genetic interaction analysis either with *er-105* or *erl1-2; erl2-1*. (C-G) Expression patterns of EPFL2. Histone H2B fused TdTomato was used as a reporter. Developing pistil in stage 8 flower (C, D), and stage 9 flower (E, F). (G) A magnified view of a white box in (E). Transverse cross section (D, F) were obtained along the lines. v and r indicate valve and replum, respectively. Bar = 100 µm. Student’s t-test was used for the statistical analyses (A) and Tukey-Kramer’s test was used for the statistical analyses (C, D). Different letters indicate significant difference (p< 0.005).

### Loss of *EPFL2* causes irregular patterning and twinning of ovules

We observed abnormal ovules and seeds in both *epfl2* mutants. In some cases (0.27%), two ovules were initiated and developed from a single funiculus (Figure 5A, B). Although EPFL2 functions in the ERL1 and ERL2 pathway, this twin-ovule phenotype was not observed in *erl1-2; erl2-1* plants, but in the *epfl2-2; erl1-2; erl2-1* and *epfl2-2; er-105*, indicating that EPFL2 can act also independently of ER-family receptors. In *epfl2* mutant plants, neighboring cells on the placenta appear to acquire ovule identity and differentiate into ovules, resulting in twins. In order to visualize early ovule initiation patterns, we introduced *pDORNRÖSCHEN(DRN)::erGFP* as a marker for the earliest stages of ovule initiation. During embryogenesis, *MP* activates expression of the auxin-responsive transcription factor *DRN* in the tip of cotyledons^37, 38^. The semi-quantitative auxin reporter R2D2^39^ revealed that auxin maxima are established at the tip of ovule primordia coinciding with *DRN* expression (Figure 5C-E), indicating that *DRN* expression also reflects auxin distribution^37, 38^, and thus can serve as marker to visualize ovule initiation patterns. Before ovule initiation, *DRN* was ubiquitously expressed in the placenta (Figure 5F, G), but when placental cells acquire ovule identity, *DRN* expression becomes confined to the ovule initial cells (Figure 5H, I). In wild-type plants, the ovules initiated with 2 to 4 cells intervals (Figure 5J, L). In *epfl2-2* mutants, *DRN* was expressed in a much broader pattern (Figure 5K) and *DRN* expression domains appeared less regularly spaced (Figure 5K, L). We quantified the spacing by counting the number of cells between adjacent ovule primordia. In the wildtype, we found on average 2.97 cells between two ovule initial cells, and these average values were slightly increased for the *epfl2* mutant lines (3.10 cells). Importantly, cell numbers in *epfl2* varied from 1 to 6 cells, whereas the wild-type displayed a very regular ovule spacing with cell numbers ranging between 2 to 4 (Figure 5L). We conclude that EPFL2 serves the initiation of ovules at regularly spaced intervals.

**Figure 5.**
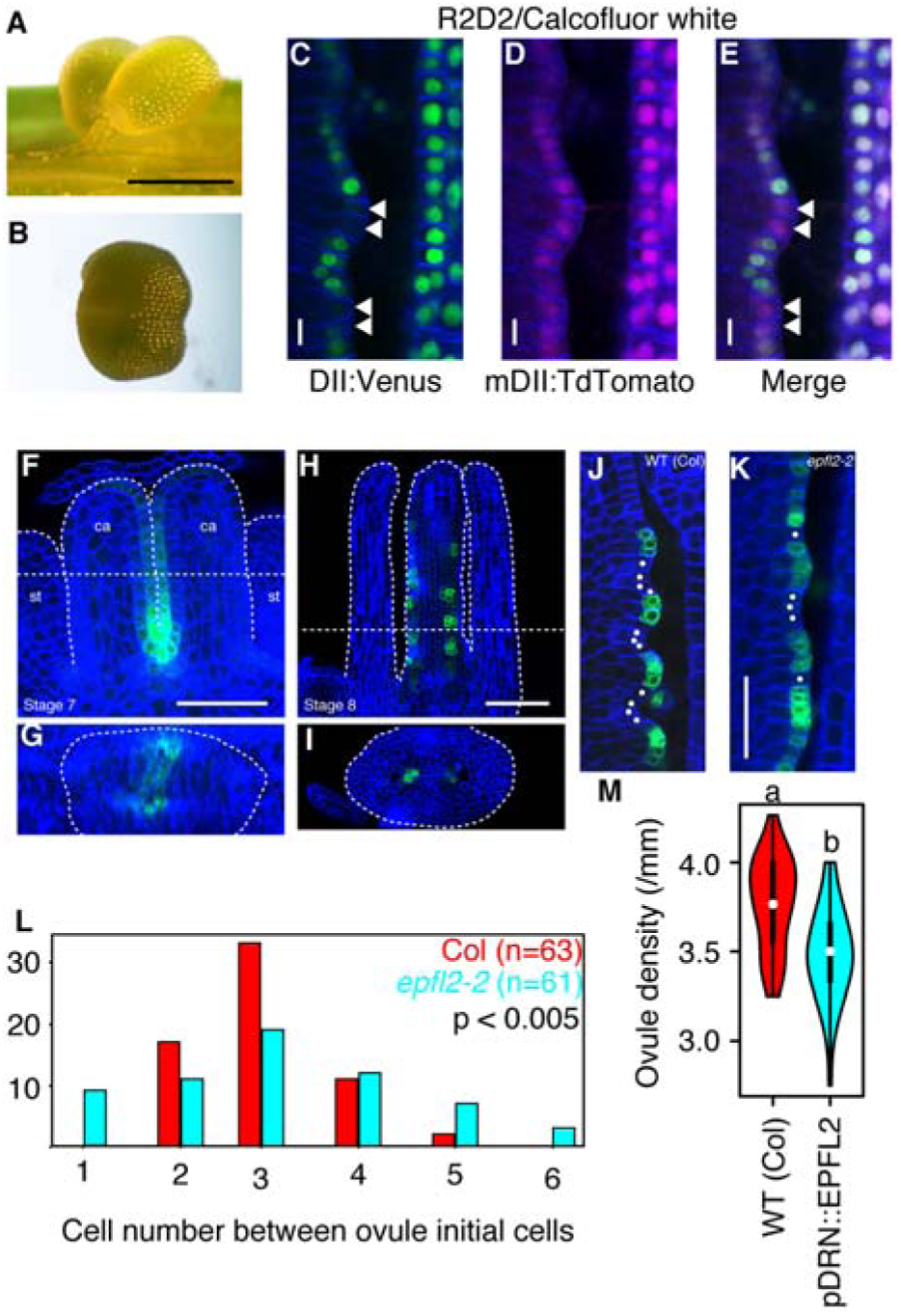
Disrupted regular patterning in ovule spacing in *epfl2* and ectopic expression of EPFL2. (A, B) Ovule twinning phenotype in *epfl2-2*. (C-E) R2D2 expression in developing ovule primordia. Arrow heads indicate auxin maxima. (F, G) Expression pattern of *DRN* before acquiring ovule identity. (H, I) Expression pattern of *DRN* after arising ovule primordia. (J) Initiation of ovule primordia in wild-type (*pDRN::GFP*; Col) plants and (K) *epfl2* mutant (*pDRN::GFP*; *epfl2-2*) plants. (L) Quantification of cell number between ovule initial cells. Counted cells were indicated by dots in (J) and (K) as an example. F-test was used for the statistical analysis. (M) Seed density (seed number/fruit length (mm)) phenotype in *pDRN::EPFL2*; Col transgenic plants. Student’s t-test was used for the statistical analysis. Different letters indicate significant difference (p< 0.005). Scale bar = 500 µm (A, B), 50 µm (C, D, E, F, H), 20 µm (J, K).

### ER-family receptors are coexpressed with EPFL2 and EPFL9 in pistils

From our genetic analysis, we concluded that two major pathways control ovule patterning. One is the EPFL9/ER pathway that mainly controls fruit growth, the other is the EPFL2/ERL1/ERL2 pathway which has a major impact on ovule density via regulating ovule initiation patterns, and also weakly contributes to fruit growth. We next analyzed the expression profiles of *ER*, *ERL1*, *ERL2*, *EPFL9* and *EPFL2*. ER-family receptors were previously shown to be expressed in different parts of the pistil^15, 40^. For the analysis of ER, ERL1 and ERL2, we used translational fusion lines with YFP as a reporter^24, 41^ and for EPFL9 and EPFL2 we generated transcriptional reporter lines using EGFP and TdTomato as fluorescent tags with Histone H2B^17, 21^. To visualize the expression patterns, we combined tissue clearing^42^ and confocal microscopy. In stage 8 flowers, ER was broadly expressed in various organs including carpels (Figure 2D), consistent with previous observations^43^. In the pistils, ER was mainly expressed in the valve (Figure 2E)^43^, but signal was also weakly detected in ovule primordia and inter-ovule spaces (Figure 2E). ERL1 expression was not detected in the carpels at early stages of development (Figure 2F). ERL2 was expressed in the carpels including the placenta before ovule primordia became apparent (Figure 2H). When ovule primordia were initiated, the expression of ERL1 and ERL2 was detected in inter-ovule spaces and the ovule primordia (Figure 2G, I). The signal of ERL2 was strongly visible at the boundary and the tip of ovule primordia which will develop into the nucellus, the outer and the inner integuments, but signal was weaker in the basal domian of ovule primordia (Figure 2I). Compared to ERL2, ERL1 signals were weaker and somehow patchy (Figure 2G). ERL1 and ERL2 were only weakly expressed in valves (Figure 2G, I). As expected from the STOMAGEN RNAi phenotype, EPFL9 was exclusively expressed in the inner cell layers of the valves (Figure 3D, E) from stage 8 onwards (Figure 3B, C) but lacking at the valve margin and the replum (Figure 3E). The expession patterns are consistent with EPFL9 acting as a short range signal that controls fruit growth via ER. In contrast to EPFL9, EPFL2 expression was detected in the placenta (Figure 4E, F), and importantly, once the ovule primordia were initiated, confined to the inter-ovule spaces (Figure 4E, F, G). In transverse sections, EPFL2 expression was also visible in the valve, around the valve margin and the replum (Figure 4F). However, expression was not noted in carpels of stage 8 flowers (Figure 4C, D). As previously reported^44^, EPFL2 seems to be preferentially expressed at the boundary between ovules.

### Altered expression of EPFL2 affects the ovule initiation pattern

The *DRN* expression profile in the placenta is largely complementary to that of *EPFL2* (Figure 4C, D, E, F, G, 5F, G, H, I). To further test the importance of *EPFL2* in the pattern of ovule initiation, we characterized the transgenic plants which misexpress *EPFL2* from the *DRN* promoter (Figure 5F, G, H, I). When *EPFL2* misexpression was driven by the *DRN* promoter in a wild-type background (*pDRN::EPFL2),* the resulting transgenic plants carried fruit with a length similar to the non-transgenic siblings (Figure S9), but showed a significantly reduced ovule number and ovule density (Figure 5M, Figure S9). This indicated that EPFL2 cannot promote fruit growth from the placenta domain, and that a regular and interspersed expression of EPFL2 is required for proper spacing of ovule initiation.

### Quantitative analysis reveals interactions between peptides and receptors

Our genetic analysis indicates that EPFL2 acts preferentially via ERL1 and ERL2; however, earlier co-immunoprecipitation experiments in *Nicotiana benthamiana* have shown that EPFL2 can physically associate with all ER-family receptors *in vivo*^17^. Because EPF/EPFL peptides may bind to ER-family receptors with different affinities^45^, we investigated interaction properties between ER-family receptors and their ligands EPFL9 and EPFL2 *in vitro*. Recombinant peptides and the extracellular domains of ER-family receptors were expressed in *E. coli* and purified. Because EPFL2 and EPFL9 are cysteine-rich and their final conformation is stabilized by the formation of specific disulfide bonds, the peptides were first affinity purified, refolded in refolding buffer, and separated from unfolded peptides by HPLC^24^. We then evaluated peptide bioactivity by measuring their impact on stomatal density. As a control, six cysteine residues of EPFL2 were substituted to serine residues (EPFL2 (CS)), which should render the peptide less stable. As observed in transgenic EPFL2-overexpressing Arabidopsis lines^17, 35^, EPFL2 treatment reduced the number of stomata (Figure S11B, I), whereas EPFL9 had the opposite effect (Figure S11D, I). Furthermore, since TOO MANY MOUTH (TMM) is a stomatal lineage specific co-receptor protein^46^ and not expressed at the carpel wall and the placenta (Figure S10), we also used *tmm-KO* plants. *tmm* knockout mutants were found to be sensitized for EPFL2 and responded more strongly (Figure S11F, I), which is consistent with previous studies^35, 45^. These assays indicated that the purified peptides were functional. We then tested the direct interactions between the peptides and receptors using isothermal titration calorimetry (ITC). Both EPFL2 and EPFL9 bound to the extracellular domain of ER-family receptors, albeit with different affinities. EPFL2 showed a binding preference for ERL1 and ERL2 (Figure S12). EPFL9 bound to ER, ERL1 and ERL2 with similar affinities (Figure S12).

From our combined data, we propose that regular spacing of ovules at defined intervals is coordinated with fruit growth through the EPFL2/ERL1/ERL2 and EPFL9/ER signalling pathways.

## Discussion

In selfing species such as Arabidopsis, pollen availability is no longer a limiting factor for fertilization, and the key determinant for seed production is now ovule number. The total number of seeds that can be generated by an individual plant depends on the available resources which can be invested into the formation of branches carrying flowers.

Overall reproductive success then depends on the total number of flowers, and the number of ovules that are being initiated in each individual flower. In the developing Arabidopsis ovary, ovules are initiated from the placenta, and the total length of the pistil at the time of ovule initiation restricts the maximum number of ovules that can be formed. Not suprisingly, there is a general correlation between fruit length and overall seed number, so that ecotypes that generate longer siliques also bear more seeds. The genetic basis of ovule initiation and fruit growth has been studied in much detail, and a number of key transcription factors and phytohormones have been investigated which control the formation of ovules and fruit development. However, the fundamental patterning mechanism that determines the spacing of ovule anlagen within the placenta remained unexplored.

Organ initiation in plants generally requires auxin accummulation at discrete sites. Using the auxin-regulated transcription factor DRN as a sensor for auxin signalling, we found that an evenly distributed auxin signal in the developing placenta is resolved into a regularly spaced pattern of founder cells of the young ovules. Importantly, this patterning process is not a repetitive process and takes place in a structure with a finite size, which clearly distinguishes it from other well studied patterning processes in plants, such as phyllotaxis or stomatal patterning. We therefore carried out a natural variation analysis and QTL analysis to identify genes responsible for ovule density, leading to the identification of *ER* and *EPFL2*. Additionally, we found that the ER paralogs ERL1 and ERL2, as well as the EPF/EPFL-family peptide EPFL9, also regulate ovule density. Our genetic interaction studies and expression analysis showed that these two pathways control ovule density in distinct ways. The EPFL9 pathway, acting from the carpel wall, exclusively controls fruit elongation without affecting the ovule initiation pattern. Reduction of ovule number observed in *er-105* seems to be an indirect consequence of the smaller fruit size and the limited availability of space. The EPFL2 pathway also affects fruit growth, but has a more pronounced impact on the patterning of ovule initial cells and thus increases ovule density (Figure 6A, B).

**Figure 6.**
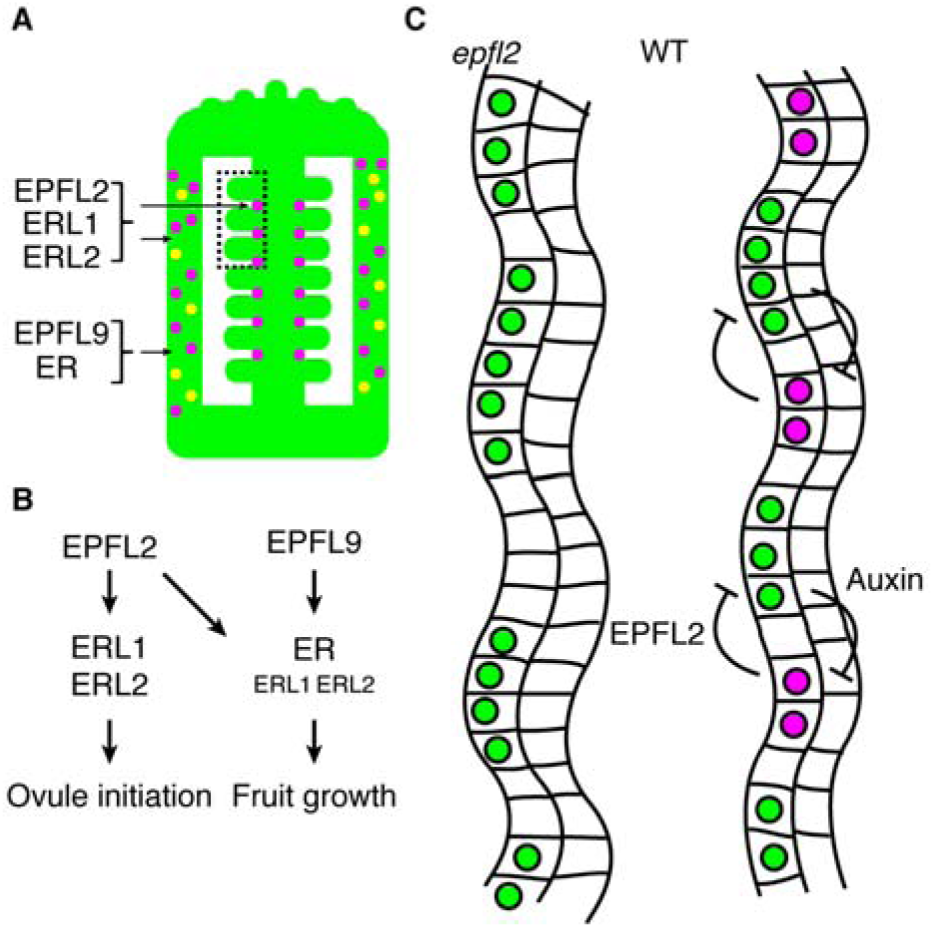
Coordination of fruit growth and ovule patterning by two peptide-receptor pairs. (A) Graphical summary of expression patterns of EPFL2 (magenta) and EPFL9 (yellow). (B) Two pathways control ovule initiation pattern and fruit growth to archive coordinated fruit growth. (C) The interaction between EPFL2 (magenta) and auxin (green) establishes regular ovule pattern in placenta.

We observed ovule twinning in *epfl2* mutant plants, which is caused by mis-patterning during ovule initiation. Furthermore, ectopic expression of EPFL2 also caused ovule patterning defects, which we interpret that EPFL2 is a dosage-sensitive regulator of ovule initiation. Our genetic, biochemical and expression data further suggest that ERL1 and ERL2 are the main receptors for EPFL2. Among these two, ERL2 has a major role as a receptor for EPFL2 for two reasons: First, the *erl2-1* mutant enhanced the *er-105* phenotype more severely than *erl1-2*, and second, ERL1 is expressed at lower levels than ERL2 in the placenta. Ovule twinning was observed in *epfl2-2; erl1-2; erl2-1* as well as *epfl2-2* mutants, but not in *erl1-2; erl2-1* mutants, suggesting that in the absence of ERL1 and ERL2, other receptors contribute to EPFL2-mediated ovule initiation. TMM is a well-characterized co-receptor for ER-family receptors in stomata development, but since TMM is not expressed in the pistil (Figure S10) ^46^, it is not likely to act here.

The transcriptional regulation of EPFL2 is so far unknown, but it is tempting to speculate that the transcription factors, CUC1, CUC2, and CUC3 are potential upstream regulators of *EPFL2* expression, because their expression profiles are similar to that of EPFL2 ^10, 47^, and *CUC1 RNAi; cuc2* plants produce fewer ovules than wild-type plants ^10^. Transcriptomic analysis revealed that some of EPFL family genes were downregulated in *CUC1 RNAi; cuc2* plants^7^. *cuc2-3; cuc3-105* mutants can carry twinned ovules ^47^, as seen in *epfl2*, and *EPFL2* expression under the control of the *CUC2* promoter was sufficient to restore the leaf margin serration phenotype of *epfl2* mutants ^17^.

Prior to ovule primordia initiation, *DRN* expression was uniformly observed at the placenta (Figure 5C). Since *DRN* is a direct target of MP ^38^, auxin seems to be signalling uniformly during early stages of placenta growth. However, once placenta cells adopt ovule identity, auxin maxima are established at the tip of ovule initials ^2^ (Figure 5C-E) and *DRN* expression is confined to these positions (Figure 5H). We propose that similar to the feedback mechanism that operates in leaf margin development ^17^, *EPFL2* restricts auxin accumulation to the developing ovule primordia, while auxin at the same time provides a feedback signal that supresses *EPFL2* expression (Figure 6C). This auxin-*EPFL2* negative feedback loop provides an important new element to control precise and highly regular ovule spacing patterns, which safeguards equal nutrient access as a bet hedging strategy. With its additional function in promoting fruit growth due to expression in the carpel wall, *EPFL2* thus serves to integrate these two processes. Life history variations that necessitate trade-offs between seed number, seed size and fruit size could then act through differential expression of *EPFL2*.

Cultivated asian rice was selected for awnlessness to facilitate harvesting. One of the underlying causal mutation inactivates an *EPFL2*-orthologue, OsEPFL1, which also affects grain length and grain number, giving rise to more compact panicles with more seeds ^48–50^. Although the underlying mechanism in rice is not yet understood, it highlights a role of *EPFL2*-genes as evolutionary conserved integrators of ovule initiation patterns, seed number, seed size and floral organ development ^51^.

## Materials and Methods

### Plant materials and growth conditions

For natural variation analysis, 96 *A. thaliana* ecotypes ^52^ were planted and grown in continuous light at either 16°C or 21°C. After germination, the plants were vernalized for 6 weeks at 4°C. For QTL analysis, 165 RILs ^5^ were planted and grown in continuous light at 16°C. For phenotypic and expression analysis, plants were grown under long-day conditions (16-h photoperiod). The Van-0, Van-0 *ER+*, Hir-1, and Hir-1 *ER+* accessions were previously described ^30^. The L. *er; epfl2-1* (CSHL_ET5721), L. *er; epfl2-1; ER+,* and L. *er; ER+* lines were previously described ^17^. An *epfl2* mutant in the Columbia accession was generated by CRISPR/Cas9. For this, we designed two single guide RNAs targeting *epfl2* (Figure S6A) in one vector. We obtained five different *epfl2* mutant alleles in the T2 generation (Figure S6B), all of which caused a smoother leaf margin phenotype indicative of *epfl2*-mutants (Figure S6C) as previously reported ^17^. For further analysis, #36-45 which has a genomic deletion between the sgRNA1 and 2 target sites was selected. Hereafter we refer to the original *epfl2* mutant in L. *er* (CSHL_ET5721) as *epfl2-1* and our new allele #36-45 as *epfl2-2*. Consistent with our previous analysis using *epfl2-1*, we found that *epfl2-2* phenocopies *epfl2-1* (Figure 4A, Figure S7), and that *epfl2-2* mutants also resemble the *erl1-2; erl2-1* phenotype (Figure 2A and 4A). The *gER:YFP; er-105*, *gERL1:YFP; erl1-2* and *gERL2:YFP* lines were previously described ^24, 41^. The *er-105, erl1-2, erl2-1, er-105; erl1-2, er-105; erl2-1, erl1-2; erl2-1* lines were previously described ^15^. The STOMAGEN RNAi line was generated ^21^. The *pDRN::GFP; Col* was described ^38^. The auxin semi-quantitative marker line R2D2 was developed in ^39^. The *epfl2-2* and *epfl2-3* lines were generated by CRISPR/Cas9 genome editing in this study, and the *epfl2-2; er-105, epfl2-2; erl1-2; erl2-1* were generated by genetic crossing. We also generated the following transgenic plant lines: *pEPFL2::H2B:TdTomato;* Col*, pEPFL9::H2B:EGFP:3HA:His; Col, pDRN::EPFL2;* Col, and *pDRN::GFP; epfl2-2*. Plants were transformed using *Agrobacterium tumefaciens* strain GV3101 or C58 pSOUP via the floral dip method.

### QTL analysis

For the RIL population, the final phenotype value for each line was calculated as the average of all the replicates. The Genotype information from 243 markers in the Ler/Cvi RIL map was collected from available published data ^5^. QTL analysis was performed within the R statistical software with the qtl package ^53^ using a Multiple QTL Mapping (MQM) approach. In the MQM mapping approach, we used a forward stepwise approach preselecting the ERECTA marker as a cofactor.

### Plasmid constructs

The plasmids and primers used in this study are listed in Table S1. Vectors pFH1 and pFH6 ^54^ along with the in-house vector pUB-Cas9-@EPFL2 were used for the knockout of *EPFL2* with the CRISPR/Cas9 system. To generate the construct pEPFL2::H2B:TdTomato, *EPFL2* promoter DNA was amplified by PCR from Col genomic DNA and inserted at the HindIII and SmaI sites of pPZP211/35S using the InFusion kit (Clontech) yielding the intermediate vector pPZP211/pEPFL2. The *H2B* (At5g22880) gene was amplified from Col cDNA and was inserted into the SmaI and SacI sites of pPZP211/pEPFL2 using the same method, yielding vectors pPZP211/pEPFL2::H2B. Finally the TdTomato gene was amplified and inserted into the SacI and SacII sites of pPZP211/pEPFL2::H2B to generate pPZP211/pEPFL2::H2B:TdTomato. To generate the construct pEPFL2::EPFL2, the *EPFL2* coding sequence was inserted into pPZP211/pEPFL2 at the BamHI and SacII sites. To construct the pDRN::EPFL2 vector, the *DRN* promoter and terminator sequences and the *EPFL2* coding sequence were amplified by PCR from Col genomic DNA or cDNA and inserted into vectors pGGA000, pGGE000, and pGGC000 respectively. The pGGZ001, pGGA000-pDRN, pGGB002, pGGC000-EPFL2, pGGD002, pGGE000-tDRN and pGGF007 DNA fragments were then assembled by GreenGate cloning ^55^. For pEPFL9:H2B-EGFP(or TdTomato):3HA:His, the *H2B* gene was amplified as above and inserted into the SmaI and SacI sites of pPZP211/35S:EGFP(or TdTomato)-3HA-His ^56^. The plasmids were digested with XbaI and EcoRI, and the H2B-EGFP (or TdTomato) 3HA-His:NosT fragments were transferred to vector pPZP211. The *EPFL9* promoter was amplified from Col genomic DNA and inserted into the vectors at the SalI and SmaI sites using the InFusion kit as above. Codon optimized mature EPFL2 and mutated mature EPFL2 sequences were synthesized (Thermo Fisher Scientific) and cloned into the SacII and XhoI sites of pET41a. Mature EPFL9 was amplified from *Arabidopsis thaliana* Col cDNA and cloned into pGEX4T1 by Gibson assembly (NEB). Ectodomains of ER (E25-R580), ERL1 (M26-R582), ERL2 (M28-R585) were amplified from *Arabidopsis thaliana* Col cDNA and cloned into pETEV16 by Gibson assembly (NEB).

### Photography of leaves

To characterize the leaf margin, the seventh leaf of each plant was photographed under a Nikon SMZ25 stereomicroscope.

### Data visualization and statistical analysis

R (version 3.5.1) was used for data visualization and statistical analysis. The following statistical tests were used to calculate the corresponding p-values. A two-tailed Student’s t-test was used for pairwise comparisons, whereas Dunnett’s test and Tukey-Kramer’s test were used to compare multiple sets of data to a control or all possible pairs, respectively. F-test was used to compare two variations. In each case, a value of p < 0.005 was considered significant.

### Tissue clearing and expression analysis

Flowers and pistils were dissected under a standard dissection microscope. The samples were fixed in freshly prepared 4% paraformaldehyde in PBS (pH 7.4) supplemented with 0.05% Silwet L-77 for 3–5 h under vacuum, followed by incubation in ClearSee as previously described [41]. The tissues were then stained with Calcofluor White to visualize the cell walls. The processed tissues were observed under a confocal microscope (Carl Zeiss LSM780, Carl Zeiss LSM880 or Leica TCS SP8). For GFP, the excitation wavelength was 488 nm and the signal was detected at 500–550 nm. For YFP, the excitation wavelength was 514 nm and the signal was detected at 520–575 nm. For TdTomato, the excitation wavelength was 561 nm and the signal was detected at 565–600 nm. For Calcofluor White, the excitation wavelength was 405 nm and the signal was detected at 415–475 nm. These ranges were selected to avoid overlaps between the signals.

### Peptide expression, purification and refolding and protein expression

Mature EPFL2 (MEPFL2) and mutant mature EPFL2 (mMEPFL2; C60S, C65S, C68S, C71S, C119S, C121S) were heterologously expressed in *E*. *coli* BL21 (DE3) as GST-His-tagged fusion proteins. EPFL9 was expressed only as GST-fusion protein in *E. coli* BL21. Peptides were purified via GST affinity chromatography by FPLC (Äkta Prime Plus, GE Healthcare). The GST tag was proteolytically cleaved by TEV-protease digestion. Peptides were separated from free GST and residual protease via reverse phase-HPLC (Supelcosil, LC-18 HPLC column, 15×4.6 cm, 3µm particle size) under an acetonitrile gradient (0-100% v/v) with 0.1%TFA (v/v). After vacuum assisted solvent evaporation, peptide pellets were resolved in refolding buffer as previously described to introduce proper disulfide bridges, which were indirectly verified by the stomata density based bioactivity assay. Peptide identities and purities were confirmed by mass spectrometry which revealed two additional amino acids (Gly-His) were attached at N-terminus of each peptide, as results of the TEV-protease cleavage.

Receptor domains of ER (E25-R580), ERL1 (M26-R582), ERL2 (M28-R585) and TMM (F24-G475) were heterologously expressed in *E. coli* BL21 (DE3) with an N-terminal His-tag and TEV-protease target sequence. The expressed protein domains were purified via Ni^2+^ affinity chromatography by FPLC (Äkta Prime Plus, GE Healthcare). The eluted proteins were subjected to buffer change by PD10 desalting column into ITC-buffer (25 mM BisTris-HCl pH6.0, 150 mM NaCl, 50 mM L-arginine and 50 mM L-glutamic acid).

### Peptide bioassay

Col and *tmm* knockout (*tmm-KO,* Salk_011958) seeds were sterilized and sown on half-strength MS medium. Prior to germination, seeds were kept in the dark at 4°C for 3 days, then transferred to continuous light at 22°C for germination. One day after germination, the seedlings were transferred to 1 ml half-strength MS liquid medium supplemented with 5 µM of the appropriate peptide in 0.5 g/L MES-KOH (pH 5.7) and were incubated as above for 5 days. At the end of the treatment period, the cotyledons were stained with 1 µg/ml propidium iodide and observed under a Confocal microscope (Carl Zeiss LSM710, Carl Zeiss LSM880 or Leica TCS SP8). For excitation, 561 nm laser line was used and signal was collected between 565–650 nm. The MES-KOH buffer (pH 5.7) without peptides was used for the mock treatment.

### ITC

ITC-experiments were carried out in a MicroCal iTC200 (Malvern Instruments) at 25°C with a sample cell of 280 µL and an injection syringe of 40 µL. Peptide pellets were dissolved in ITC-buffer and peptide concentrations were assessed by FTIR with a DirectDetect system (Merck). Protein concentrations of the receptor domains were measured by absorption at 280 nm and calculated by their molar absorption coefficient at 280 nm. The molar coefficients for ER, ERL1 and ERL2 (42400, 41410, and 42400 M^-1^ cm^-1^, respectively) were calculated based on ExPASy ProtParam. Final protein and peptide concentrations are as indicated. For each experiment 19 injections of 2 µL with a spacing of 150s were performed.

## Acknowledgement

We thank the Center for Advanced imaging (CAi) at the Heinrich-Heine University Düsseldorf for support with microscopy. We are grateful for receiving Arabidopsis seeds from the following colleagues: Martijn van Zanten (Van-0, Van-0 ER+, Hir-1, Hir-1 ER+), Wolfgang Werr (pDRN::GFP), Dolf Weijers (R2D2), Dominique Bergmann (gERL2:YFP), Ikuko Hara-Nishimura and Tomoo Shimada (STOMAGEN RNAi). This work was supported by the DFG through the Cluster of Excellence on Plant Sciences (CEPLAS, EXC1028).

## Supplementary Figures

**Figure S1. Natural variation analysis on fruit length and seed number**

96 Arabidopsis thaliana natural accessions were analyzed on fruit length and seed number per fruit at 16°C and 21°C. Each point indicates average values of fruit length and seed number. Acccessions in red points are presented in Figure 1B.

**Figure S2. Complementation of L. er, Van-0 and Hir-1 by functional ER genomic sequence**

(A) Fruit length phenotype. (B) Seed number per fruit. (C) Seed density. Student’s t-test was used for the statistical analyses. Different letters indicate significant difference (p< 0.005).

**Figure S3. Fruit length and seed number in ER family receptor mutants**

(A) Fruit length (mm) and (B) seed number per fruit were analyzed in Columbia background mutants. Tukey-Kramer’s test was used for the statistical analyses. Different letters indicate significant difference (p< 0.005).

**Figure S4. Fruit phenotype of STOMAGEN RNAi plants**

(A) Fruit length (mm) and (B) seed number per fruit were analyzed. STOMAGEN RNAi plants were generated [17]. Student’s t-test was used for the statistical analyses. Different letters indicate significant difference (p< 0.005).

**Figure S5. Genetic interaction analysis between *er* and *epfl2* in Landsberg**

(A) Fruit length (mm) and (B) seed number per fruit were analyzed in Landsberg background mutants. L. *er ER+* was used as for wild-type plants. Tukey-Kramer’s test was used for the statistical analyses. Different letters indicate significant difference (p< 0.005).

**Figure S6. Generation of *epfl2* knockout mutant in Col background by CRISPR/Cas9**

(A) Design of sgRNA. Boxes and lines indicate exons and introns, respectively. Green and white boxes indicate coding regions corresponding to mature peptide and 3’ untranslated region (UTR), respectively. Arrowheads indicate the position of sgRNA1 and sgRNA2. (B) Sequences of newly isolated *epfl2* mutant alleles. Black bars indicate the positions of sgRNA and red bars indicate the positions of protospacer adjacent motif (PAM), respectively. (C) Leaf margin phenotype of *epfl2* mutant alleles. Seventh true leaves were photographed according to [13]. Scale bars = 1 cm.

**Figure S7. Fruit phenotype of *epfl2* mutants**

(A) Fruit length (mm) and (B) seed number per fruit were analyzed. Student’s t-test was used for the statistical analyses. Different letters indicate significant difference (p< 0.005).

**Figure S8. Genetic analysis of *epfl2* and ER family receptors: Fruit length and seed number phenotypes**

Double mutant analysis of *epfl2-2; er-105* in fruit length (A) and seed number per fruit (B). Triple mutant analysis of *epfl2-2; erl1-2; erl2-1* in fruit length (C) and seed number per fruit (D). Tukey-Kramer’s test was used for the statistical analyses. Different letters indicate significant difference (p< 0.005).

**Figure S9. Fruit phenotype of *pDRN::EPFL2* plants**

(A) Fruit length (mm) and (B) seed number per fruit were analysed. Student’s t-test was used for the statistical analyses. Different letters indicate significant difference (p<0.005).

**Figure S10. Expression pattern of *TMM***

pTMM::GUS expression in stage 5-10 flower buds (A), stage 11 flower (B) and stage 12 flower (C). Scale bar = 2 mm.

**Figure S11. Evaluation of recombinant peptide bioactivity based on stomatal density**

Representative images of epidermis after peptides and mock treatments in wild-type (A-D) and *tmm-KO* mutant (E-H). Scale bar = 100 µm. (I) Quantification of stomatal number per 0.2 mm^2^. Each treatment were compared to mock treatment. Dunnett’s test was used for the statistical analyses. A value of p < 0.005 was considered as significant.

**Figure S12. Quantitative interaction analyses between EPFLs and ER-family receptors by using ITC**

Isothermal titration calorimetry of the ERECTA family receptor domains with EPFL2 (A-C) and EPFL9 (D-F). 18 injections of 2µL of peptide (50 µM) were titrated into 280 µL of the receptor domain (5 µM) at a stirring rate of 750 rpm. The experiment was performed at 25°C. The thermograms show the detected peaks of the heat change after each injection (upper panel), the integrated values were subjected to either the “one binding site” (A-E) fitting algorithm of the Microcal ITC-ORIGIN software, or the “two binding sites” algorithm (F). Each square represents the integrated value of the corresponding peak and the line resembles the yielded fitting curve after chi² minimization (lower panel). The table lists the calculated *Kd* values for each interaction with the theoretical stoichiometry (G).

**Supplemental Movie 1. Expression pattern of EPFL2 using two photon microscopy.**

A z series of stage 9-10 pistil expressing H2B:TdTomato under the control of EPFL2 promoter. Images were acquired with 1 µm intervals by using two photon microscopy (Nikon A1R) with 1000 nm excitation.

**Supplemental Movie 2. Expression pattern of EPFL2 using ClearSee and confocal microscopy.**

A z series of ClearSee treated stage 9-10 pistil expressing H2B:TdTomato under the control of EPFL2 promoter. Images were acquired with 1 µm intervals by LSM880 with 561 nm excitation.

