## Supplemental Figures for "A peptide pair coordinates regular ovule initiation patterns with seed number and fruit size"

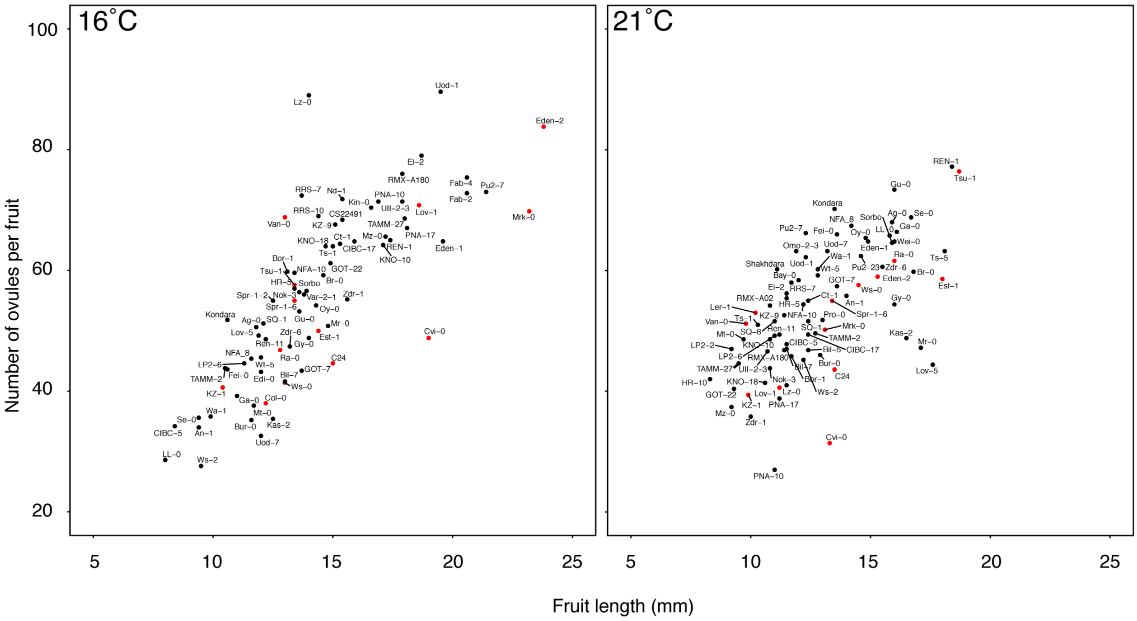


**Figure S1. Natural variation analysis on fruit length and seed number**

96 Arabidopsis thaliana natural accessions were analyzed on fruit length and seed number per fruit at 16˚C and 21˚C. Each point indicates average values of fruit length and seed number. Acccessions in red points are presented in Figure 1B.

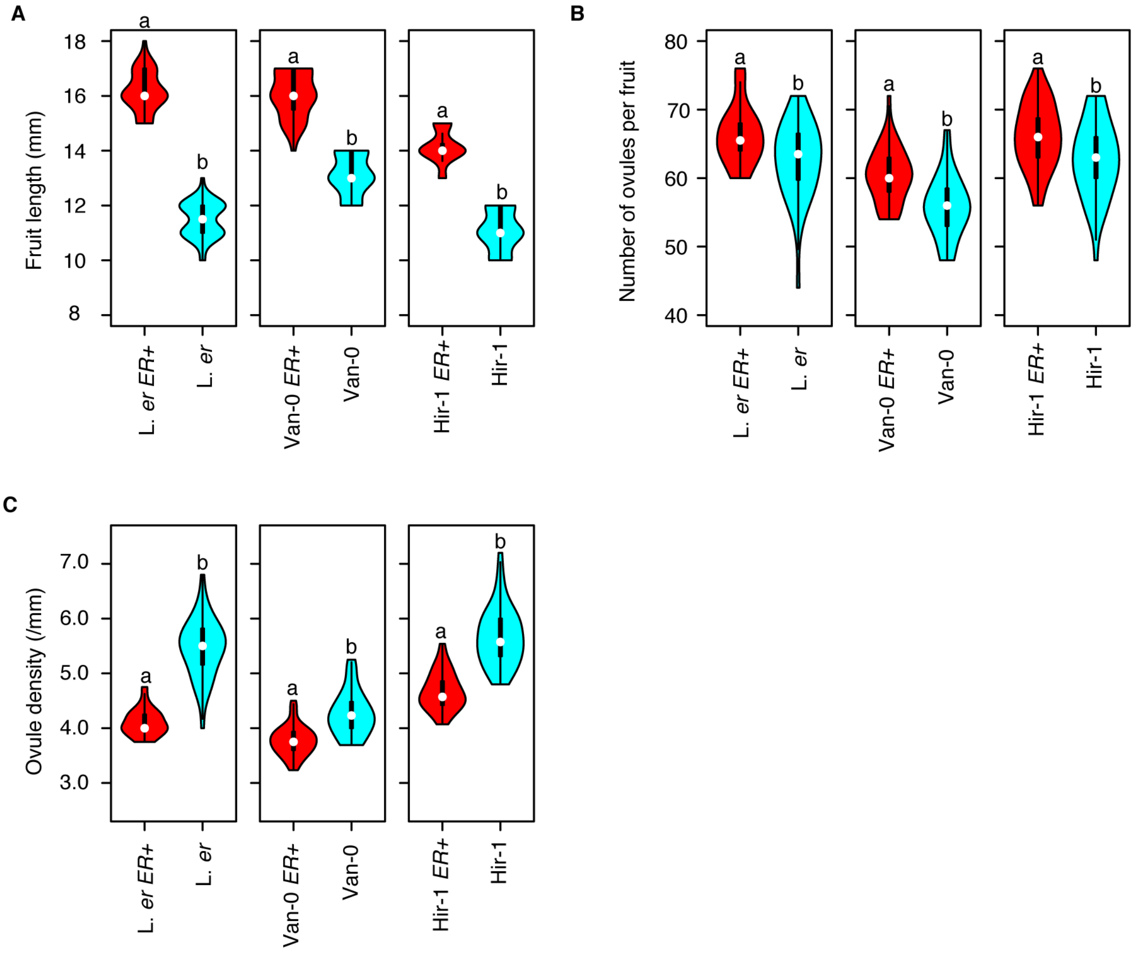


**Figure S2. Complementation of L. er, Van-0 and Hir-1 by functional ER genomic sequence**

(A) Fruit length phenotype. (B) Seed number per fruit. (C) Seed density. Student's t-test was used for the statistical analyses. Different letters indicate significant difference ($p< 0.005$).

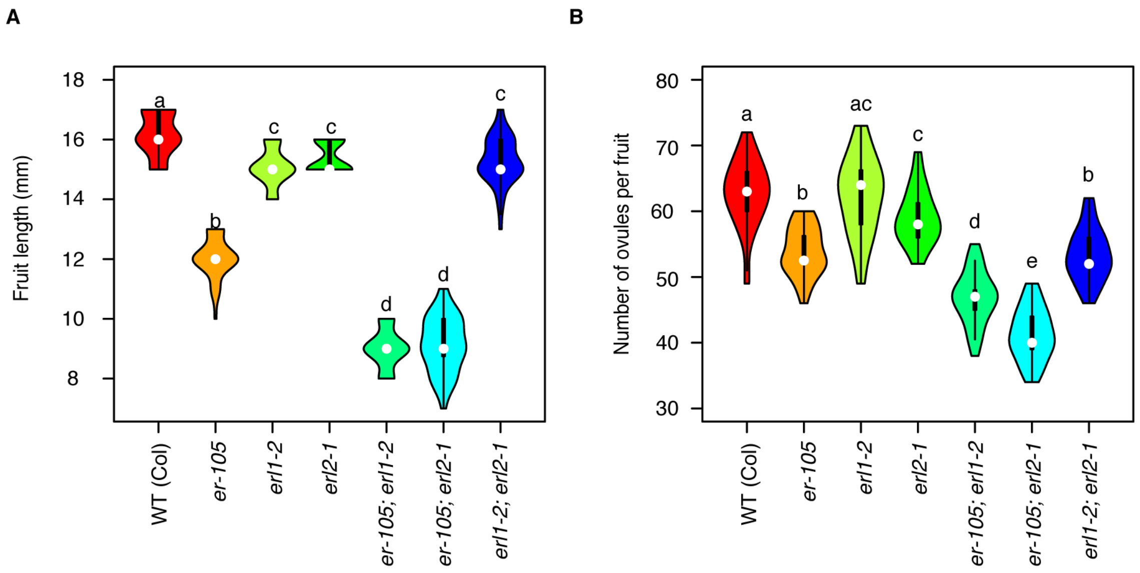


**Figure S3. Fruit length and seed number in ER family receptor mutants**

(A) Fruit length (mm) and (B) seed number per fruit were analyzed in Columbia background mutants. Tukey-Kramer's test was used for the statistical analyses. Different letters indicate significant difference ($p < 0.005$).

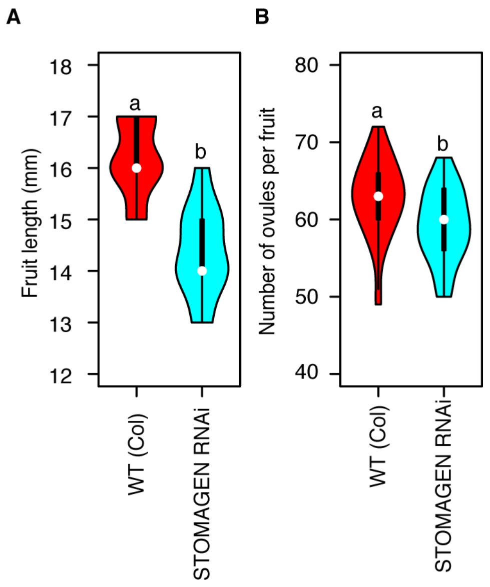


**Figure S4. Fruit phenotype of STOMAGEN RNAi plants**

(A) Fruit length (mm) and (B) seed number per fruit were analyzed. STOMAGEN RNAi plants were generated [17]. Student's t-test was used for the statistical analyses. Different letters indicate significant difference ($p < 0.005$).

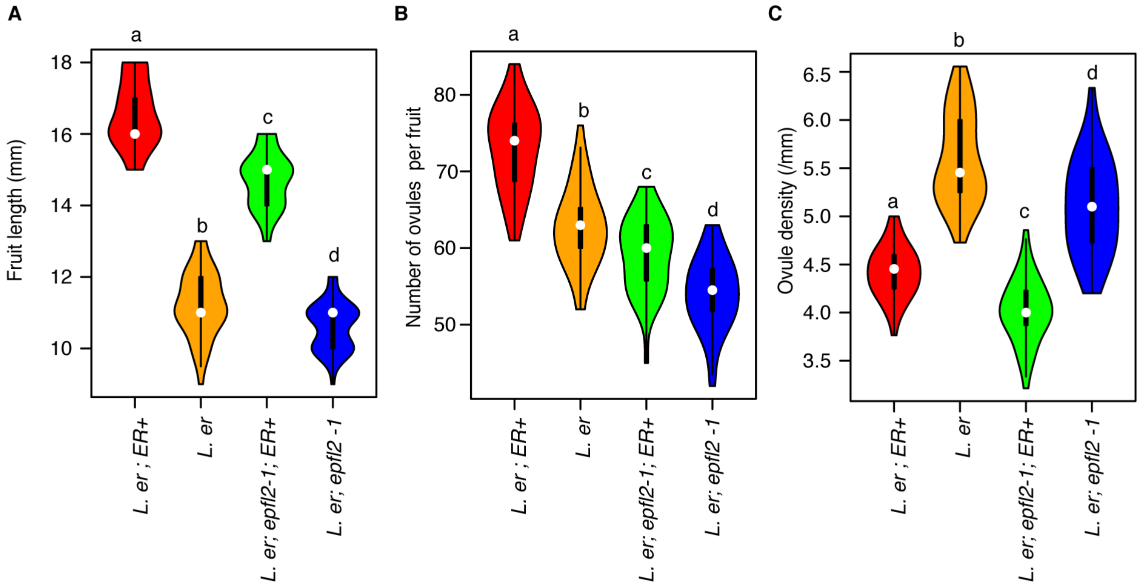


**Figure S5. Genetic interaction analysis btween *er* and *epfl2* in Landsberg**

(A) Fruit length (mm) and (B) seed number per fruit were analyzed in Landsberg background mutants. L. *er ER*+ was used as for wild-type plants. Tukey-Kramer's test was used for the statistical analyses. Different letters indicate significant difference ($p < 0.005$).

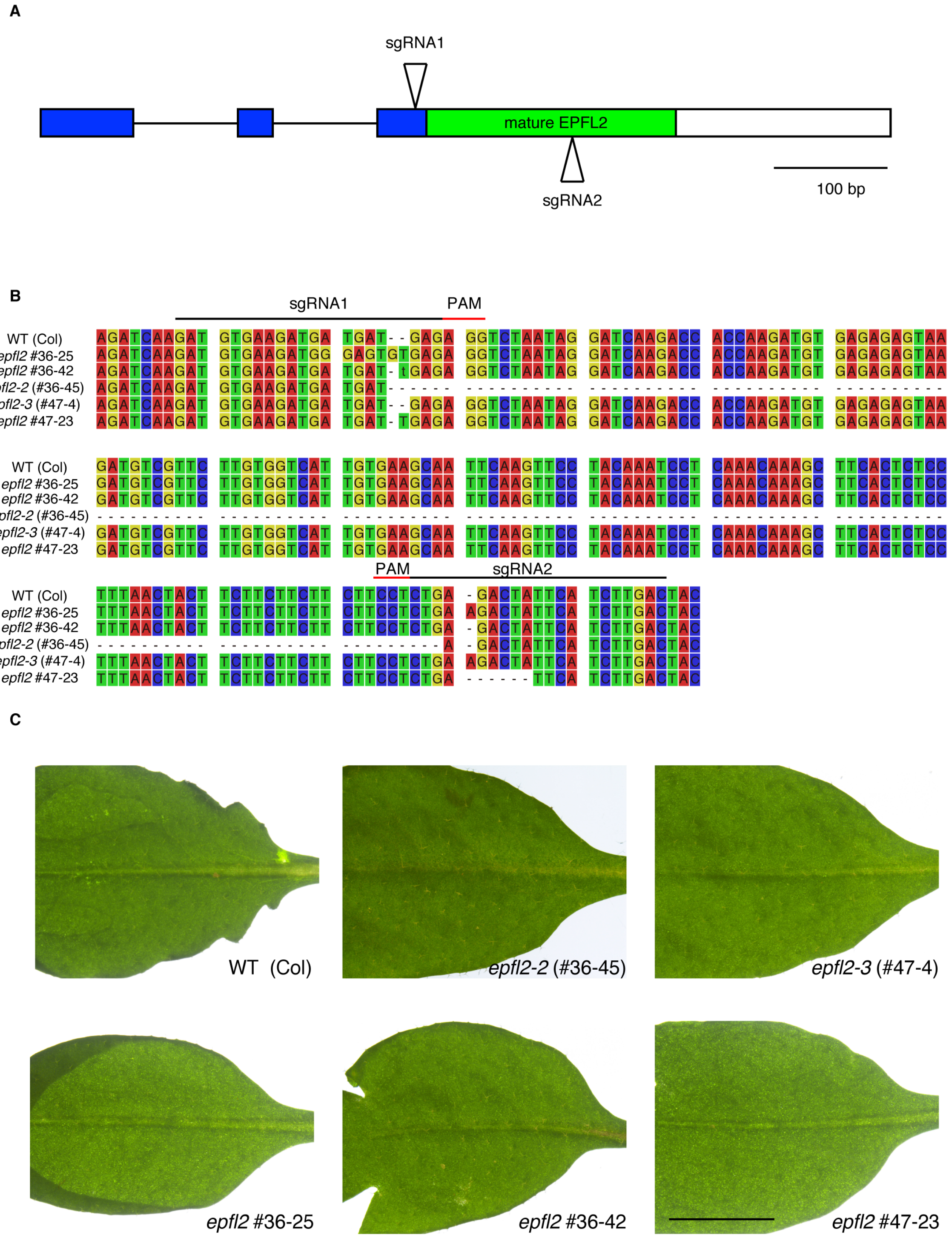


**Figure S6. Generation of *epfl2* knockout mutant in Col background by CRISPR/Cas9**

(A) Design of sgRNA. Boxes and lines indicate exons and introns, respectively. Green and white boxes indicate coding regions corresponding to mature peptide and 3' untranslated region (UTR), respectively. Arrowheads indicate the position of sgRNA1 and sgRNA2. (B) Sequences of newly isolated *epfl2* mutant alleles. Black bars indicate the positions of sgRNA and red bars indicate the positions of protospacer adjacent motif (PAM), respectively. (C) Leaf margin phenotype of *epfl2* mutant alleles. Seventh true leaves were photographed according to [13]. Scale bars = 1 cm.

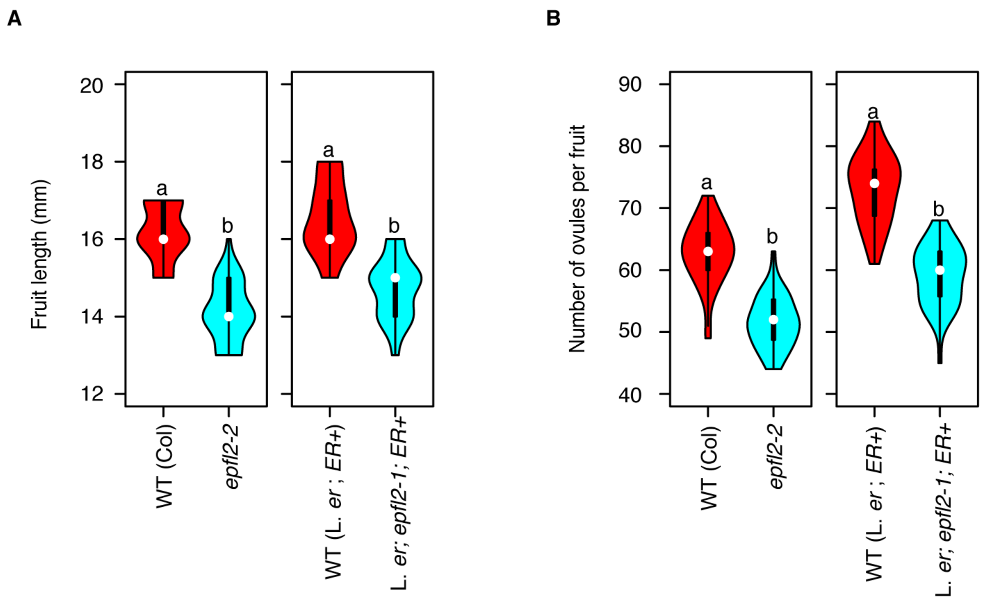


**Figure S7. Fruit phenotype of *epfl2* mutants**

(A) Fruit length (mm) and (B) seed number per fruit were analyzed. Student's t-test was used for the statistical analyses. Different letters indicate significant difference ($p < 0.005$).

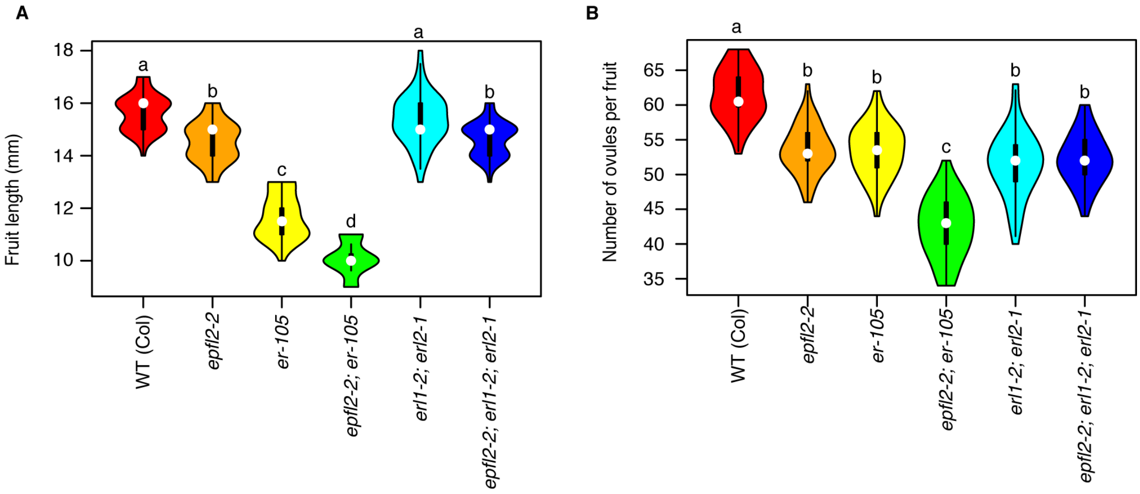


**Figure S8. Genetic analysis of *epfl2* and ER family receptors: Fruit length and seed number phenotypes**

Double mutant analysis of *epfl2-2; er-105* in fruit length (A) and seed number per fruit (B). Triple mutant analysis of *epfl2-2; erl1-2; erl2-1* in fruit length (C) and seed number per fruit (D). Tukey-Kramer's test was used for the statistical analyses. Different letters indicate significant difference ($p < 0.005$).

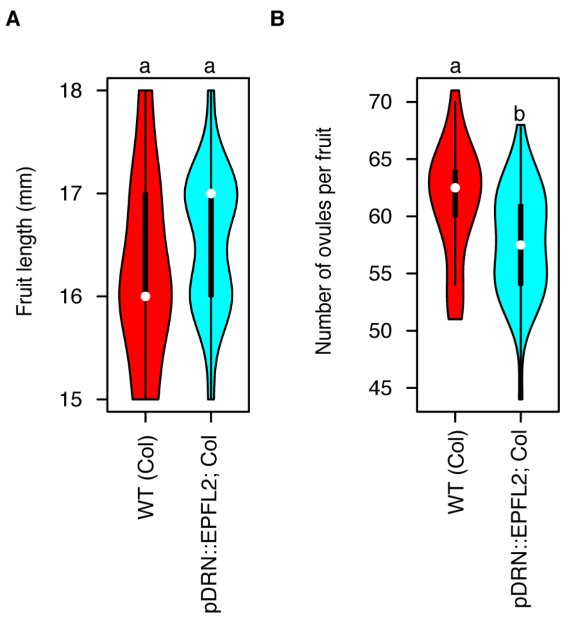


**Figure S9. Fruit phenotype of *pDRN::EPFL2* plants**

(A) Fruit length (mm) and (B) seed number per fruit were analysed. Student's t-test was used for the statistical analyses. Different letters indicate significant difference ($p < 0.005$).

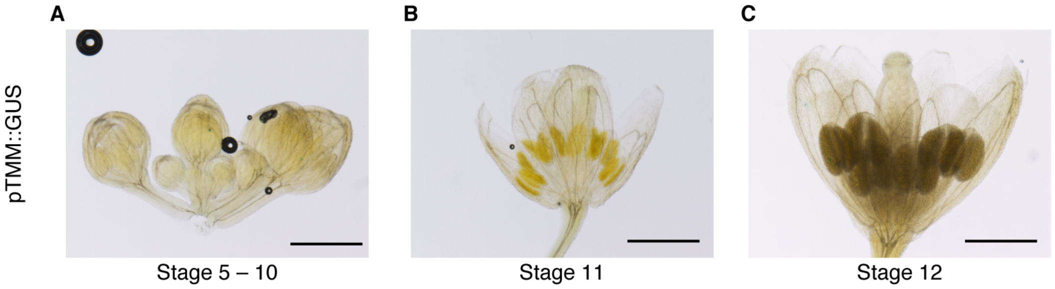


**Figure S10. Expression pattern of *TMM***

pTMM::GUS expression in stage 5-10 flower buds (A), stage 11 flower (B) and stage 12 flower (C). Scale bar = 2 mm.

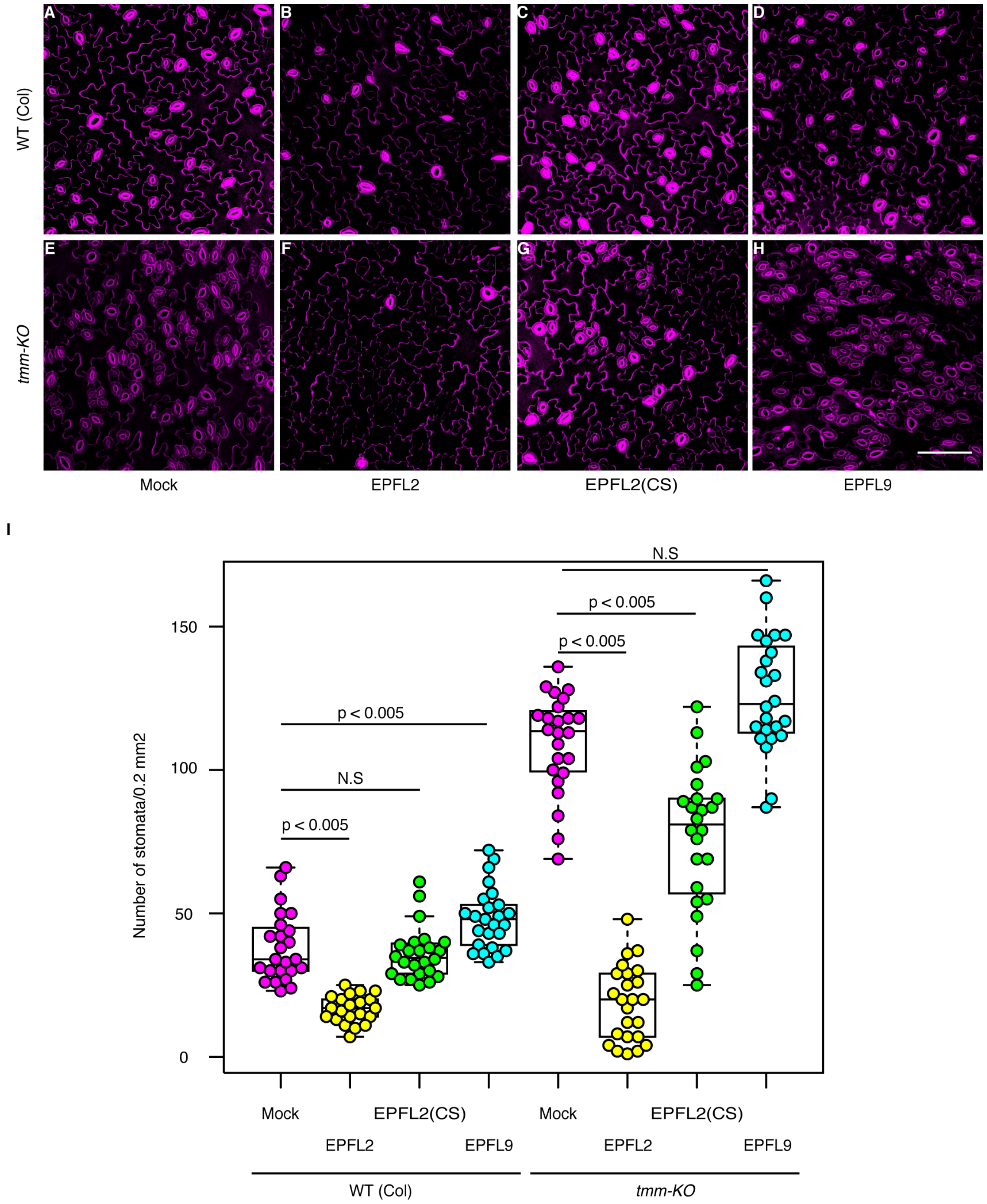


**Figure S11. Evaluation of recombinant peptide bioactivity based on stomatal density**

Representative images of epidermis after peptides and mock treatments in wild-type (A-D) and *tmm-KO* mutant (E-H). Scale bar = 100 µm. (I) Quantification of stomatal number per 0.2 $mm^2$. Each treatment were compared to mock treatment. Dunnett's test was used for the statistical analyses. A value of $p < 0.005$ was considered as significant.

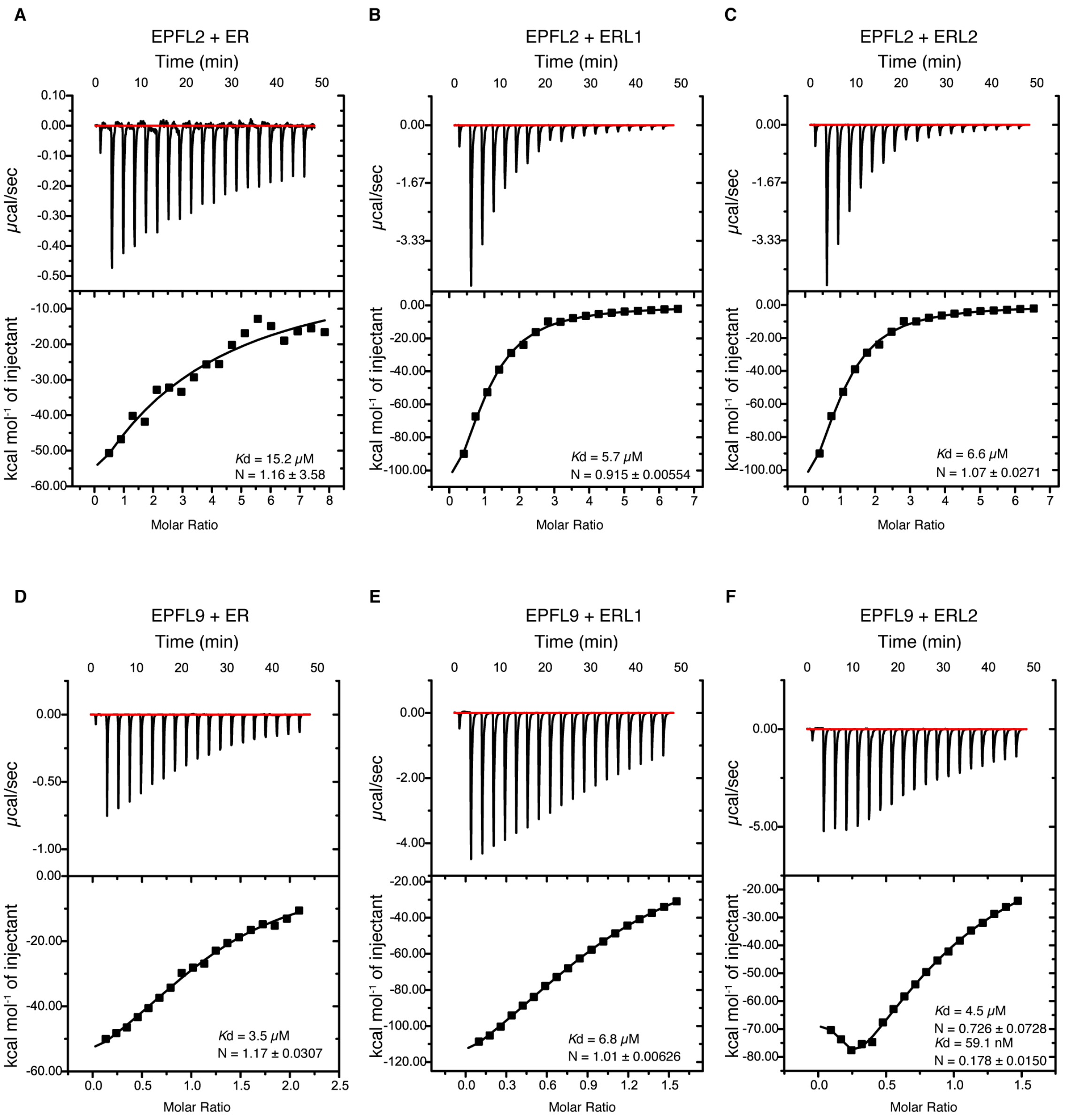


**G**

| Ligand | Receptor | *K*d | N |
|---|---|---|---|
| EPFL2 | ER | 15.2 μM | 1.16 ± 3.58 |
| EPFL2 | ERL1 | 5.7 μM | 0.915 ± 0.00554 |
| EPFL2 | ERL2 | 6.6 μM | 1.07 ± 0.0271 |
| EPFL9 | ER | 3.5 μM | 1.17 ± 0.0307 |
| EPFL9 | ERL1 | 6.8 μM | 1.01 ± 0.00626 |
| EPFL9 | ERL2 | 4.5 μM<br>59.1 nM | 0.726 ± 0.0728<br>0.178 ± 0.0150 |

**Figure S12. Quantitative interaction analyses between EPFLs and ER-family receptors by using ITC**

Isothermal titration calorimetry of the ERECTA family receptor domains with EPFL2 (A-C) and EPFL9 (D-F). 18 injections of 2μL of peptide (50 μM) were titrated into 280 μL of the receptor domain (5 μM) at a stirring rate of 750 rpm. The experiment was performed at 25°C. The thermograms show the detected peaks of the heat change after each injection (upper panel), the integrated values were subjected to either the "one binding site" (A-E) fitting algorithm of the Microcal ITC-ORIGIN software, or the "two binding sites" algorithm (F). Each square represents the integrated value of the corresponding peak and the line resembles the yielded fitting curve after $chi^2$ minimization (lower panel). The table lists the calculated *Kd* values for each interaction with the theoretical stoichiometry (G).

**Supplemental Movie 1. Expression pattern of EPFL2 using two photon microscopy.**

A z series of stage 9-10 pistil expressing H2B:TdTomato under the control of EPFL2 promoter. Images were acquired with 1 μm intervals by using two photon microscopy (Nikon A1R) with 980 nm excitation.

**Supplemental Movie 2. Expression pattern of EPFL2 using ClearSee and confocal microscopy.**

A z series of ClearSee treated stage 9-10 pistil expressing H2B:TdTomato under the control of EPFL2 promoter. Images were acquired with 1 μm intervals by LSM880 with 561 nm excitation.
